# Impact of non-proteinogenic amino acid norvaline and proteinogenic valine misincorporation on a secondary structure of a model peptide

**DOI:** 10.1101/2022.09.26.509456

**Authors:** Zara Škibola, Ita Gruić Sovulj, Aleksandra Maršavelski

**Affiliations:** Department of Biology, Faculty of Science, University of Zagreb, Zagreb 10000, Croatia; Department of Chemistry, Faculty of Science, University of Zagreb, Zagreb 10000, Croatia

## Abstract

Norvaline is a straight-chain, hydrophobic, non-proteinogenic amino acid, isomeric with valine. Both amino acids can be misincorporated into proteins at isoleucine positions by isoleucyl-tRNA synthetase when the mechanisms of translation fidelity are impaired. Our previous study showed that proteome-wide substitution of isoleucine with norvaline resulted in higher toxicity in comparison to the proteome-wide substitution of isoleucine with valine. Although mistranslated proteins/peptides are considered to have non-native structures responsible for their toxicity, the observed difference in protein stability between norvaline and valine misincorporation has not yet been fully understood. To examine the observed effect, we choose the model peptide with three isoleucines in the native structure, introduced selected amino acids at isoleucine positions and applied molecular-dynamics simulations at different temperatures. The obtained results showed that norvaline has the highest destructive effect on the β-sheet structure and suggest that the higher toxicity of norvaline over valine is predominantly due to the misincorporation within the β-sheet secondary elements.

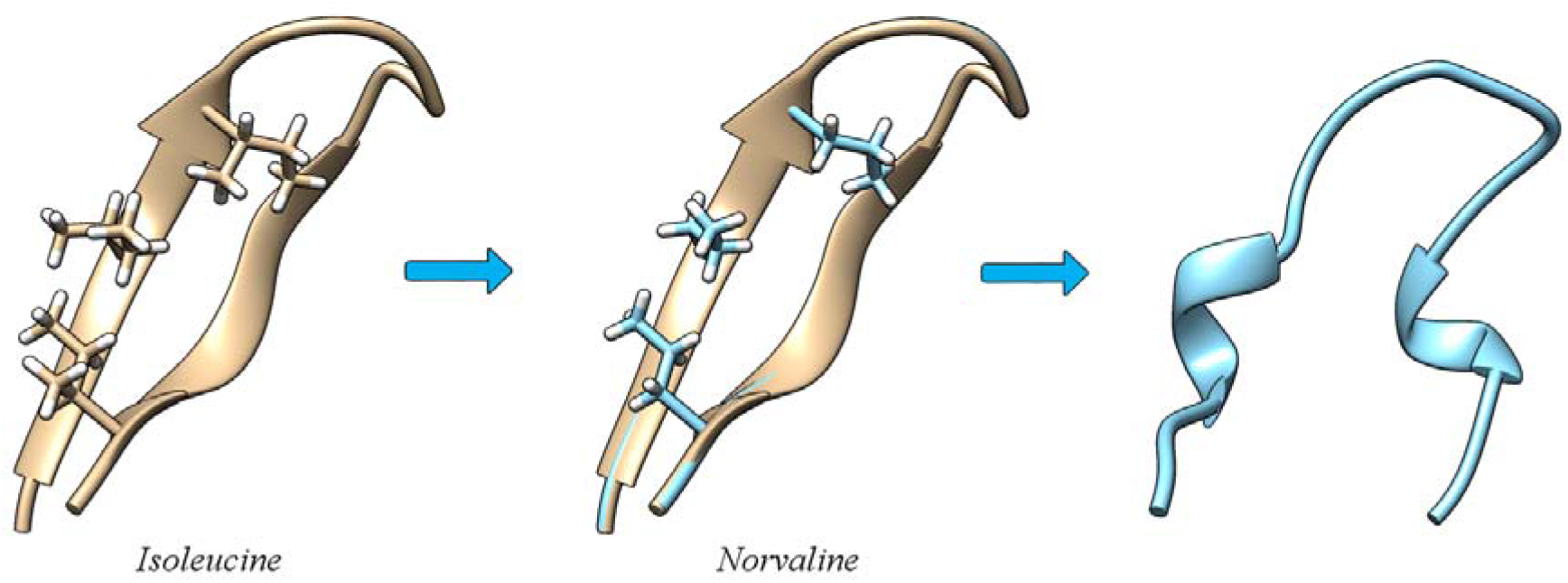

## INTRODUCTION

Norvaline (Nva) is a natural, non-proteinogenic, unbranched amino acid, not encoded by the genetic code. Nva is isomeric with proteinogenic amino acid valine (Val) and can compete with Val for misactivation by isoleucyl-tRNA synthetase (IleRS). Subsequently both amino acids could end up misincorporated into proteins at Ile positions. Although IleRS uses both pre-transfer editing of Val-AMP within the synthetic site^1^ and deacylation of misacylated Val-tRNA^Ile 2,3^ and Nva-tRNA^Ile 4^ within the post-transfer editing domain to maintain the accuracy of protein synthesis, it has been shown that in circumstances of impaired editing both Val and Nva are toxic for cell.^4,5^ Our previous study compared the effect of Nva and Val mistranslation into *E. coli* proteome at Ile positions and analysis showed that proteome-wide substitution of Ile with Nva resulted in higher toxicity in comparison to the proteome-wide substitution of Ile with Val.^4^ These three amino acids differ in their propensities towards secondary structure elements.^6,7^ Namely Ile and Val exhibit high β-sheet propensities, whereas Nva belongs to a good helix-forming residue, therefore exhibiting different effects on proteome stability upon misincorporation. Previous experimental study,^8^ where authors used combinatorial mutagenesis to examine the effects of introducing alanine, a good α-helix-forming residue and valine, a poor helix-forming residue at 12 positions in a native α-helix 1 of the N-terminal domain of λ repressor, clearly demonstrated that Ala and Val are equally well tolerated at the majority of examined sites, although showing opposite helix-forming propensities. Surprisingly, the authors also discovered that α-helix 1 of λ repressor can accommodate up to four valines at once, a poor helix-forming residue, with retention of biological activity. On the other hand, a tolerance of a β-sheet to multiple mutations, especially in light of Val/Nva misincorporation at Ile sites, has not yet been studied. To investigate why Nva misincorporation into proteins is more deleterious compared to Val^4^ we opted for *in silico* approach. We chose the envelope HIV-1 IIIB V3 peptide^9^, a natural β-hairpin with three isoleucines and we misincorporated Nva and Val at all three Ile positions. By applying classical molecular dynamics simulations at different temperatures, we could elegantly study how β-hairpin structure stability is affected by misincorporation and explain the higher toxicity of norvaline compared to valine.

## Methodology

### Model Construction

We opted for the envelope HIV-1 peptide IIIB V3^9^, a natural β-hairpin consisting of 18 amino acids forming two antiparallel β–strands, connected by a loop (Figure 1). This simple protein structural motif has three isoleucines at positions 4, 6 and 17 (Figure 1.) and by replacing isoleucines, with alanines, valines, norvalines and leucines, respectively, we obtained the models (Table 1) whose stability was tested at two temperatures 27 °C and 77 °C using classical molecular dynamics simulations. Selected amino acids differ in secondary structure propensities where Ala, Leu and Nva exhibit α-helix propensity whereas Val exhibits β–sheet propensity.^6,7^

**Figure 1.**
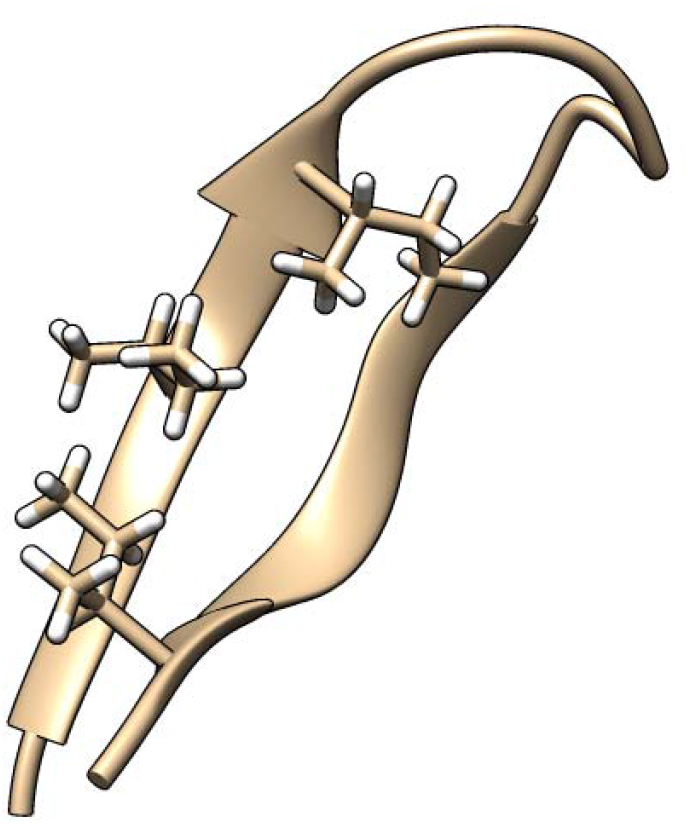
Structure of the HIV-1 IIIB V3 β-hairpin (PDB ID 1B03) with three isoleucines at positions 4, 6 and 17, all given in licorice representation. Isoleucines 4 and 6 are part of the first β–strand, whereas isoleucine 17 is part of the second β–strand.

**Table 1.**
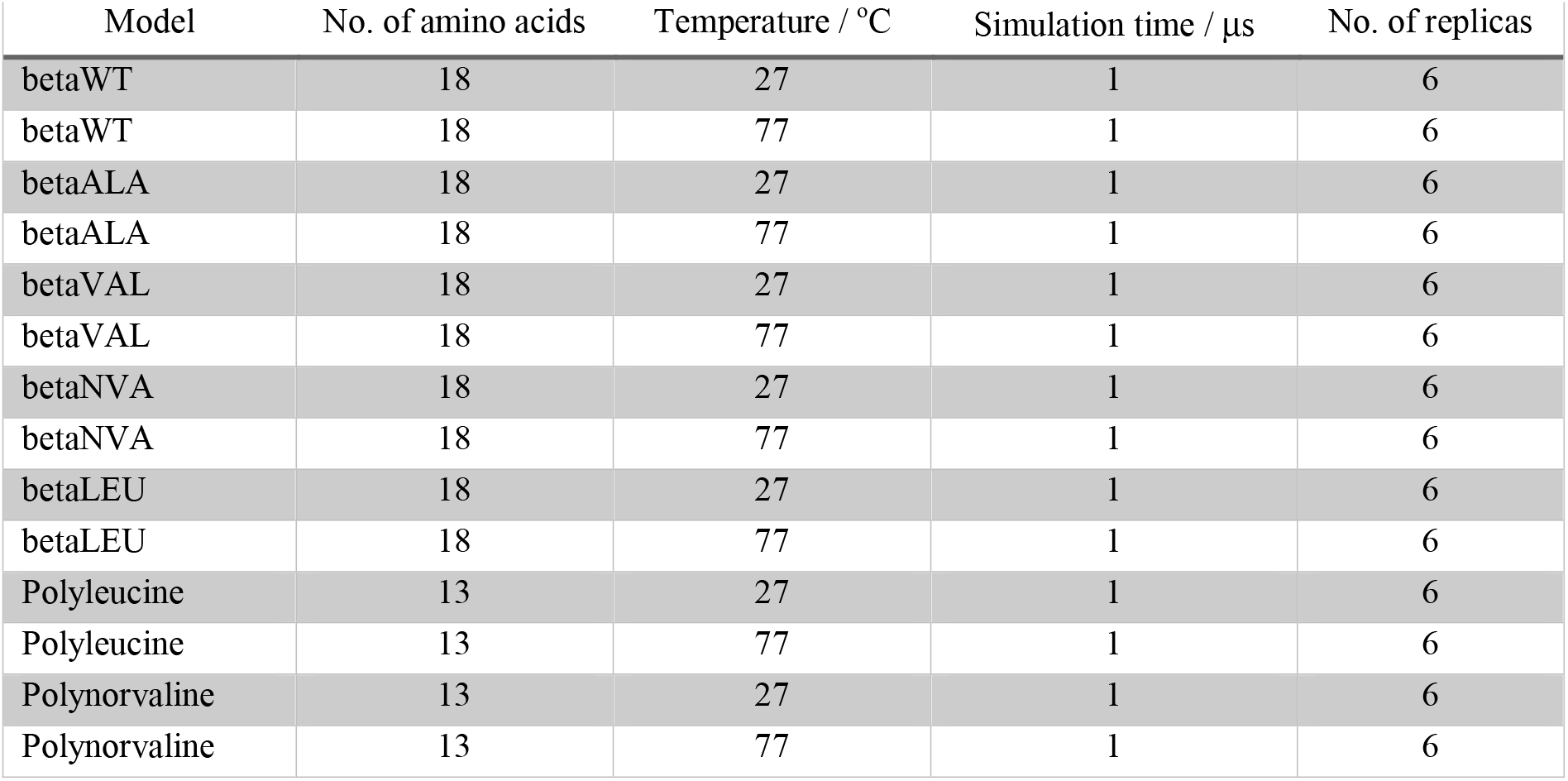
Models prepared for MD simulations. Variants of HIV-1 IIIB V3 peptide (betaWT) were obtained by changing all 3 isoleucines with corresponding amino acid.

The structure of the HIV-1 IIIB V3 peptide was taken from a protein database (PDB ID 1B03)^9^, and protein variants were designed using the Chimera^10^. The most probable rotamer was selected from the rotamer library^11^. Norvaline was built manually from a wild-type structure, where the methyl group on the C atom of each isoleucine was replaced by a hydrogen atom. Models of the HIV-1 IIIB V3 peptide of the HIV envelope were simulated at two temperatures 27 °C and 77 °C, and each model was simulated for 1 μs, in six replicas (Table 1). As can be seen in Table 1 we prepared 7 models, giving 84 simulations in total (Table 1). To further corroborate propensities for secondary structures, in light of obtained results, we also constructed linear, unfolded structures of the homopolypeptides made of leucines and norvalines using the tleap module within the AmberTools18 software package^12^. We expected that both amino acids have a propensity towards α-helix, as already known from the literature.^6,7^ Each of the homopolypeptides consisted of 13 amino acids that had acetyl (ACE) and N-methyl amide group (NME) at their N- and C-terminus, respectively. It is known that the α-helix is a macrodipole and that a negatively charged amino acid such as Asp at the N-terminus and a positively charged amino acid such as Lys at the C-terminus of the α-helix contribute to its stability. Since the models used are constructed from the same non-polar amino acids, which cannot contribute to the stabilization in this way, the ACE and NMA groups were placed at the N-end and C-terminus of the α-helix, respectively.^13^ In this way the electrostatic attraction of the charged carboxyl and terminal amino group located at the C- and N-terminus of the models is avoided.

The number of amino acids has also been shown to affect the folding of homopolypeptides into the α-helix. The helices can be divided into two groups depending on the number of amino acids that build them. Polypeptides with 6, 7, 10, 11, 13, 14, 17, 18, 21, 22, 24, 25, 28, 29 and 31 amino acids favor the helix, and those with 8, 9, 12, 15, 16, 19, 20, 23, 26, 27 and 30 amino acids are classified as less favorable for the secondary structure of the α-helix.^14^ It has been shown that the balance between entropy-driven unfolding and the folding driven by minimizing the nonpolar regions exposed to water, determines which α-helix lengths are favored^15^. These results show the critical polypeptide length with a maximum probability of forming an α-helix is that of 13 amino acids^15^ and that is why we opted for homopolypeptides built of 13 amino acids. We run molecular dynamics simulations of all prepared models at two temperatures of 27 °C and 77 °C and to credibly investigate the potential energy surface and obtain statistically relevant results each model was simulated in six replicas. An overview of the prepared models is given in Table 1.

### Molecular dynamics simulations

The parameters for norvaline were prepared *de novo* (Figure S1), whereas standard amino acids were described using the Amber ff14SB force field^16^. All constructed models were dissolved in an explicit TIP3P water model^17^ and all models were placed in a truncated octahedron-shaped simulation box. The systems were minimized in several steps and then equilibrated at the corresponding temperature (27 °C and 77 °C) at which the production runs were performed after equilibration. Each model was simulated in 6 replicas, and each replica was 1 μs and each trajectory consisted of a million structures. The simulation time of microseconds was chosen because it was previously shown that the folding time for the α-helix consisting of 30 amino acids at temperatures between 25 °C and 27 °C is in the range of 100 to 200 nanoseconds^18^. Therefore, the chosen time scale allows monitoring of the folding, but also the stability of the folded structure. Other experimental results ^19,20^, as well as results of MD simulations^21^, showed that the time required for folding of homopolypeptide consisted of 15 to 30 amino acids, ranging from 100 to 500 ns. The SHAKE algorithm was applied^22^ and a 2-fs step was used for numerical integration. The temperature was maintained using Langevin dynamics and pressure was controlled by Berendsen barostat^23^. The cutoff value was set to 9 Å. Production runs were performed with the AMBER18 on GPU using the pmemd.CUDA engine.^24–26^

### Data Analysis

The analysis was performed using the cpptraj module^27^ within the AmberTools18 package^12^. The two-dimensional structural landscape, as a measure of stability, is based on the root mean square deviation (RMSD) and the radius of gyration (Rg)^28,29^ of simulated systems at 27 °C and 77 °C. The radius of gyration and the root of the mean square deviations were calculated considering N, O, C, Cα atoms of the peptide bonds. The evolution of the secondary structure elements during the simulation time and the propensity of each amino acid in a particular structural element at a given temperature were also monitored. Elements of the secondary structure were assigned according to the DSSP classification^30^ based on the analysis of φ and ψ torsion angles and hydrogen bonds.

## Results

All six simulations of wild-type β-hairpin (betaWT) at 27 °C remained stable, in the conformation of the two antiparallel β-strands (Figure S2). At 77 °C, in two out of six replicas the emergence of the α-helix was observed in the first β-strand (Figure S3), however, it was short-lived. Simulations of the β-hairpin with three alanines (betaALA) incorporated into isoleucine positions showed that the structure was already disrupted at 27 °C. α-helix and 3_10_ helix briefly reappeared in four out of six simulations (Figure S4). At higher temperature, this destabilizing effect is even more pronounced. The structure of α-helix is detected in five replicas; however, all α-helices were short-lived (Figure S5). Simulations of β-hairpin with valine at three isoleucine positions (betaVAL) confirmed the great tendency of this amino acid towards the β-sheet secondary structure, since in all six simulations the native structure of the antiparallel β-sheet was preserved (Figure S6). At higher temperature, observed destabilization (Figure S7) was comparable to the wild-type β-hairpin at 77 °C. The introduction of norvaline into isoleucine positions in the β-hairpin (betaNVA) led to structure destabilization in two out of six replicas at 27 °C. More precisely, the first β-strand was destabilized and lost its structure (Figure S8). At 77 °C we detected significant denaturation of the β-hairpin in such a manner that α-helix appeared instead of both β-strands in all six replicas (Figure S9). Leucine in the β-hairpin (betaLEU) on average did not disrupt the structure of the antiparallel β-sheet, which was preserved in five out of six simulations at 27 °C (Figure S10). At higher temperature (77 °C) there was a significant change in the structures. All 6 simulations showed structural transformation towards α-helix in both β-strands (Figure S11), however, the stability and occupancies of the α-helices in all replicas were lower in comparison to α-helices formed at 77 °C in 6 replicas of betaNVA. The evolution of secondary structure elements along trajectories of all simulated systems suggests that the greatest impact on β-sheet destabilization at room temperature (27 °C) has alanine (Figure S4), whereas at high temperature (77 °C), norvaline and leucine (Figures S9 and S11) display the greatest destabilizing effect of β-hairpin. On the other hand, Val misincorporation didn’t result in any noticeable destabilization when compared to the wild-type structure at both studied temperatures.

To better compare the betaNVA and betaLEU simulations at 77 °C and to assess the contribution of norvaline and leucine to β-hairpin destabilization at higher temperatures we prepared two-dimensional structural landscapes based on the root mean square deviation (RMSD) and the radius of gyration (Rg) of simulated systems at 77 °C. As a reference structure to calculate the RMSD values between the backbone atoms N, O, C, Cα of the betaNVA and betaLEU β-hairpin structures along trajectories we used the experimentally solved structure of α-helix (PDB ID 1ALE)^40^ composed of the same number of amino acids as the β-hairpin. As previously stated, the structure of the β-hairpin (PDB ID 1B03) is composed of 18 amino acids forming two β-strands connected with a loop. In this way we could measure the similarity of the β-hairpin strands to the structure of α-helix, i.e. we could measure the destabilization of the β-hairpin structure upon mistranslation. The first β–strand is built from amino acids 2 to 6 and the second β-strand is built from amino acids 13 to 17. Therefore, we, calculated RMSD (with respect to α-helix; PDB ID 1ALE) for each β–strand of the β-hairpin taking into account backbone atoms N, C, O, Cα of the sequence comprised of amino acids from 2 to 6 and 13 to 17. The radius of gyration (Rg) along the trajectories was calculated with the respect to the reference structure of the β–hairpin whose both β–strands were transformed to α-helices, by taking into account backbone atoms N, C, O, Cα of the sequence comprised of amino acids from 2 to 6 and 13 to 17. In this way, we could quantitatively assess each structure of the obtained ensembles by comparing RMSD and Rg values of each β–strand to the structure of α-helix and β–hairpin whose strands transformed to α-helices, respectively. Lower RMSD and lower Rg values of each β–strand meant greater similarity to the structure of α-helix and a greater destabilizing effect on the β–sheet structure. As can be seen from twodimensional structural landscapes (Figures 2 and 3) norvaline has the greatest destabilizing effect on the structure of β–sheet at the higher temperature of 77 °C (Figure 2) when compared to leucine (Figure 3). This destabilization effect is more pronounced in the first strand of the β–hairpin (Figures 2 and 3, left panels).

**Figure 2.**
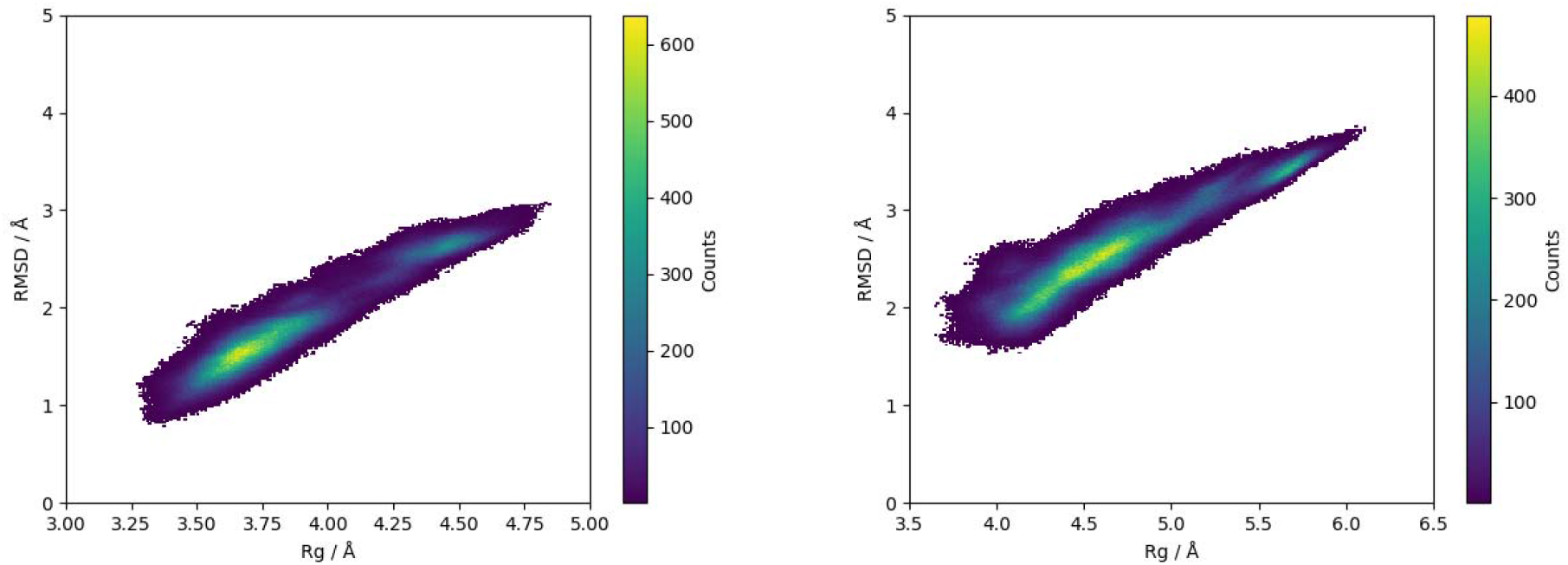
Two-dimensional structural landscapes of β-strand composed of amino acids 2 to 6 (left) and β-strand composed of amino acids 13 to 17 (right) based on the root mean square deviation (RMSD) and the radius of gyration (Rg) of **betaNVA** at 77 °C. Heat maps contain average values based on 6 independent MD runs.

**Figure 3.**
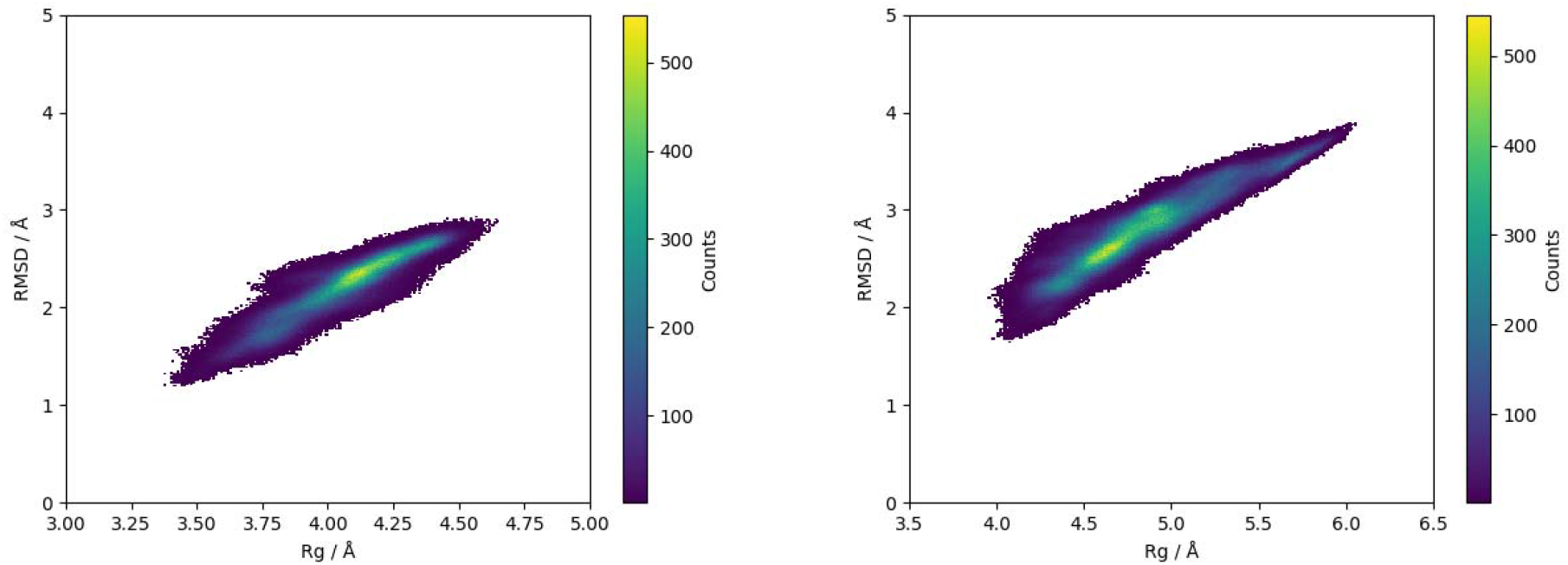
Two-dimensional structural landscapes of β-strand composed of amino acids 2 to 6 (left) and β-strand composed of amino acids 13 to 17 (right) based on the root mean square deviation (RMSD) and the radius of gyration (Rg) of **betaLEU** at 77 °C. Heat maps contain average values based on 6 independent MD runs.

We noticed that the first β-strand of betaNVA has a greater tendency towards the structure of α-helix because of having two incorporated norvalines at positions 4 and 6 in comparison to the second β-strand of betaNVA which has only one norvaline at position 17 (Figure 2).

To further assess the contribution to α-helix stability at both temperatures we analyzed folding of the linear structures composed of 13 Nva and Leu using MD simulations. Two-dimensional structural landscapes of polynorvaline and polyleucine trajectories confirmed that both amino acids have a high propensity towards α-helix since all six replicas of both structures folded into α-helix (Figures S12-S15). Heat maps (Figures 4 and 5) shows that leucine has a greater tendency toward α-helix at room temperature in comparison to norvaline. It can be seen, however, that norvaline’s tendency toward the α-helix has not been changed with the temperature (Figure 4). When it comes to leucine, the tendency to form α-helix decreases with increasing temperature (Figure 5). The obtained results showed that all structures folded on a nanosecond time scale which is expected from the literature.^31^

**Figure 4.**
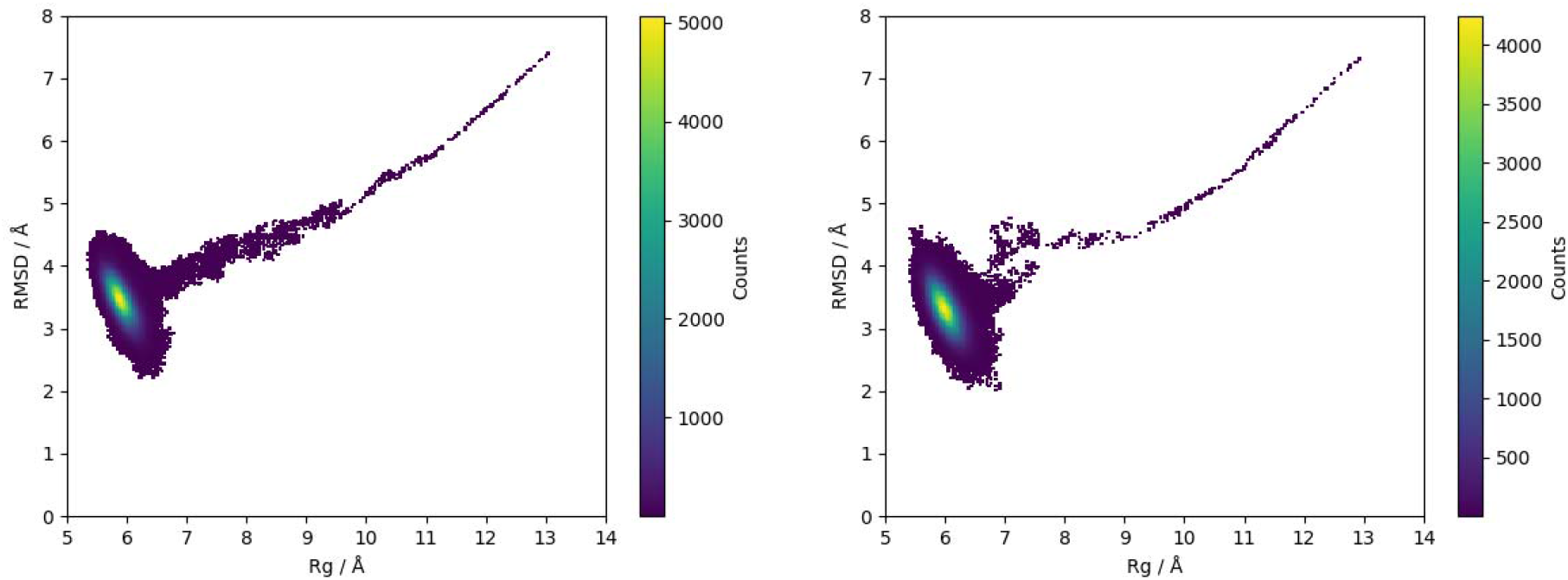
Two-dimensional structural landscapes of polynorvaline composed of 13 norvalines with ACE and NME terminal caps at 27 °C (left) and 77 °C (right) based on the root mean square deviation (RMSD) and the radius of gyration (Rg) with respect to the structure of α-helix (PDB ID 1ALE) by taking into account backbone atoms N, C, O, Cα. Heat maps contain average values based on 6 independent MD runs.

**Figure 5.**
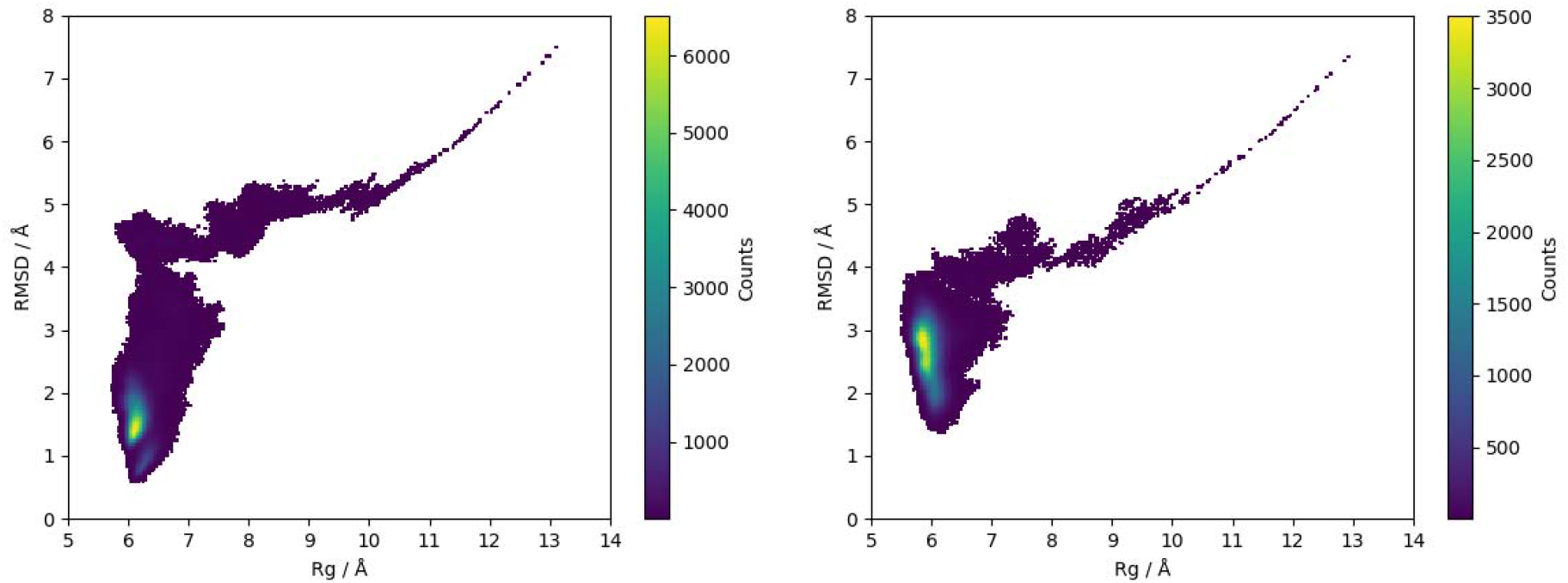
Two-dimensional structural landscapes of polyleucine composed of 13 leucines with ACE and NME terminal caps at 27 °C (left) and 77 °C (right) based on the root mean square deviation (RMSD) and the radius of gyration (Rg) with respect to the structure of α-helix (PDB ID 1ALE) by taking into account backbone atoms N, C, O, Cα. Heat maps contain average values based on 6 independent MD runs.

## Discussion and Conclusion

To understand the effect of amino acid misincorporation at isoleucine sites on protein stability, we analyzed the stability of the wild-type β-hairpin and its several variants with misincorporated Ala, Nva, Leu and Val at all three Ile positions at room (27 °C) and elevated (77 °C) temperatures. Selected amino acids differ in their propensities toward α-helix and β-sheet secondary structure elements and performed analysis showed that Nva has the highest destabilizing effect on the β-sheet structure at elevated temperature, among all simulated models. On the other hand, misincorporation of Val at Ile positions did not cause any significant destabilization of the β-hairpin structure. Additional simulations of polynorvaline folding showed interesting notion that norvaline propensity towards α-helix doesn’t change with increasing temperature. Taken altogether, Nva incorporation into β-sheet elements leads to destabilization, which is more pronounced with increasing temperature whereas α-helix structure tolerates well norvaline even at higher temperatures. These results can also be discussed in a broader sense, placing them in the context of whole protein stability. The straight alkyl side chain of Nva has greater conformational freedom than the side chains of Ile, Leu or Val, which may influence the peptide/protein folding by impacting the formation of hydrophobic core or the interaction interface between protein domains. Therefore, mistranslated proteins are prone to altered structure, in comparison to the structure of the wild-type version, loss of activity and aggregation, which all collectively affect the functionality of the proteome and viability of the cell^31^.

## ACKNOWLEDGEMENTS

A.M. would like to thank the Zagreb University Computing Centre (SRCE) for granting computational resources of the ISABELLA cluster. This work was supported by the grant from the Foundation of Croatian Academy of Sciences (10-102/384-141-2020; PI: A.M.)

## AUTHOR CONTRIBUTIONS

A.M. conceptualization of the research, conceived and designed research, data anaylsis and vizualization, supervision, wrote the original manuscript; Z.Š. performed research, data analysis and visualization, review and editing of the manuscript; I.G.-S. conceived research, review and editing of the manuscript.

