## Supplementary material for "Impact of non-proteinogenic amino acid norvaline and proteinogenic valine misincorporation on a secondary structure of a model peptide": Supporting Information-AM.docx

How non-proteinogenic amino acid norvaline affect the stability of secondary protein structures – a computational study

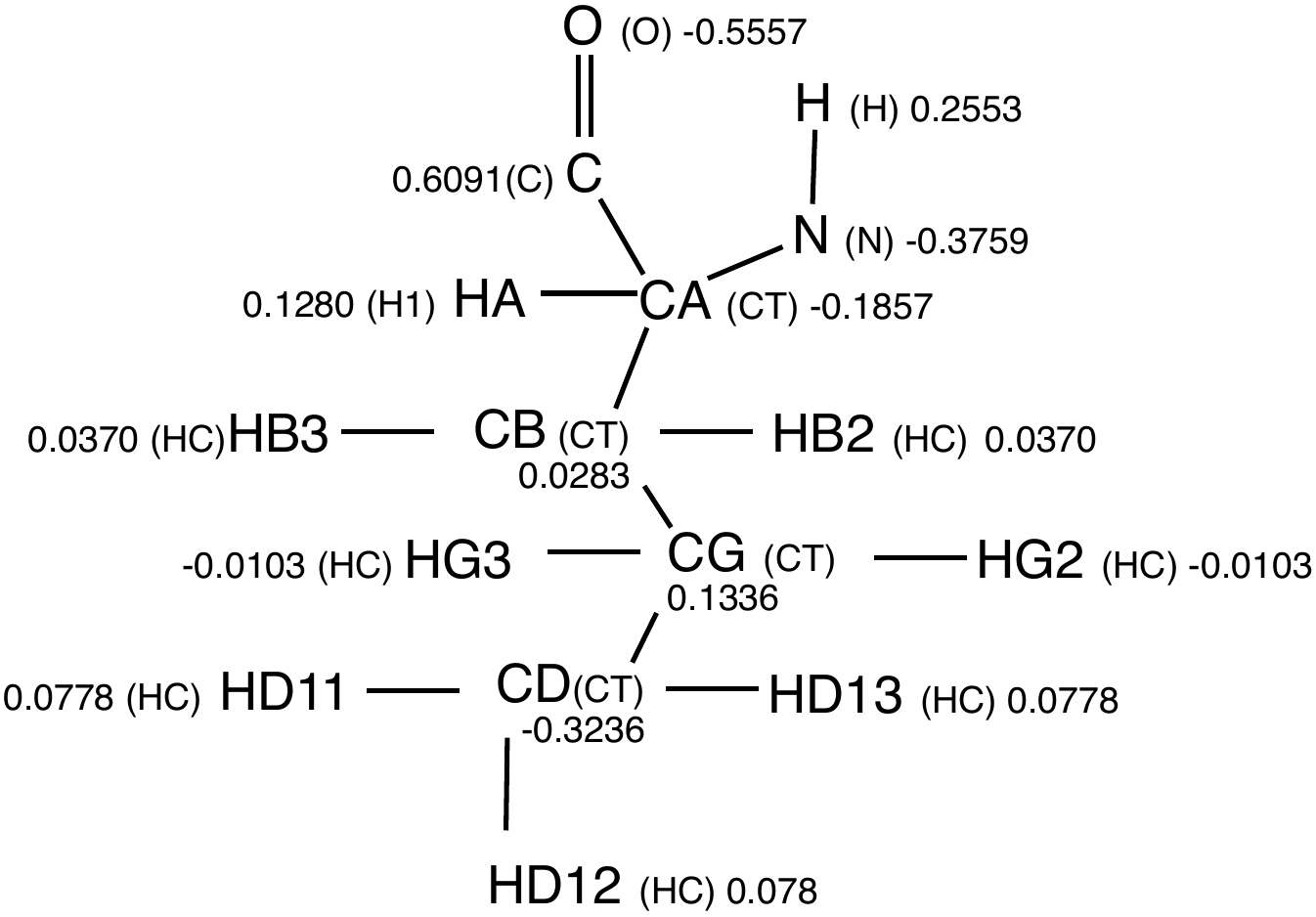

Figure S1. Parameters for norvaline with assigned atom types (given in parenthesis) and partial atomic charges.

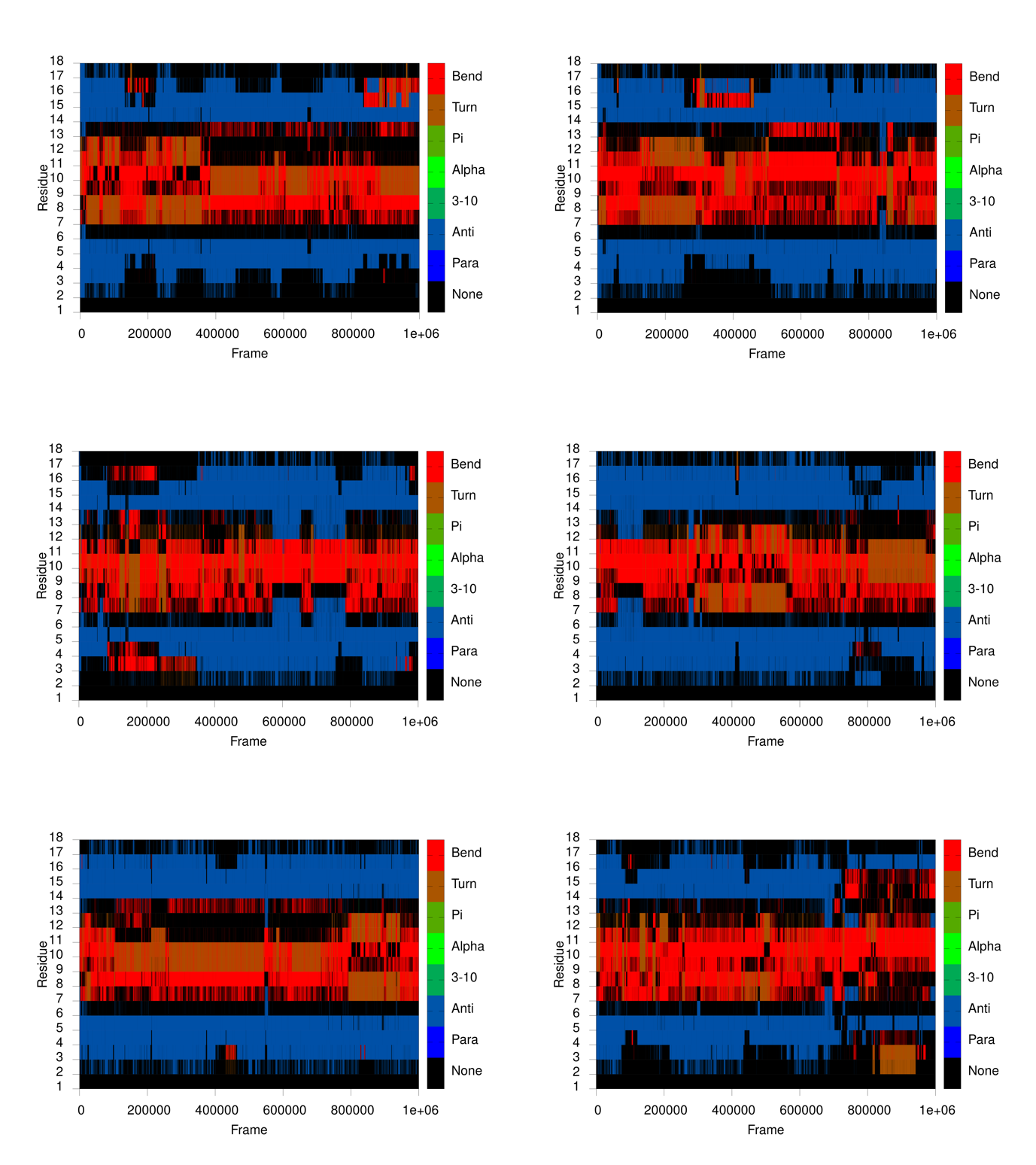

Figure S2. Evolution of secondary structure elements of HIV-1 peptide IIIB V3, wild-type β-hairpin at 27 °C during 1 μs MD simulation (six replicas). Colors denote certain element of secondary structure. Para - parallel β-sheet, Anti - antiparallel β-sheet, 3-10 – 3_10_ helix, Alpha - α-helix, Pi - π-helix, Turn - β-turn and Bend.

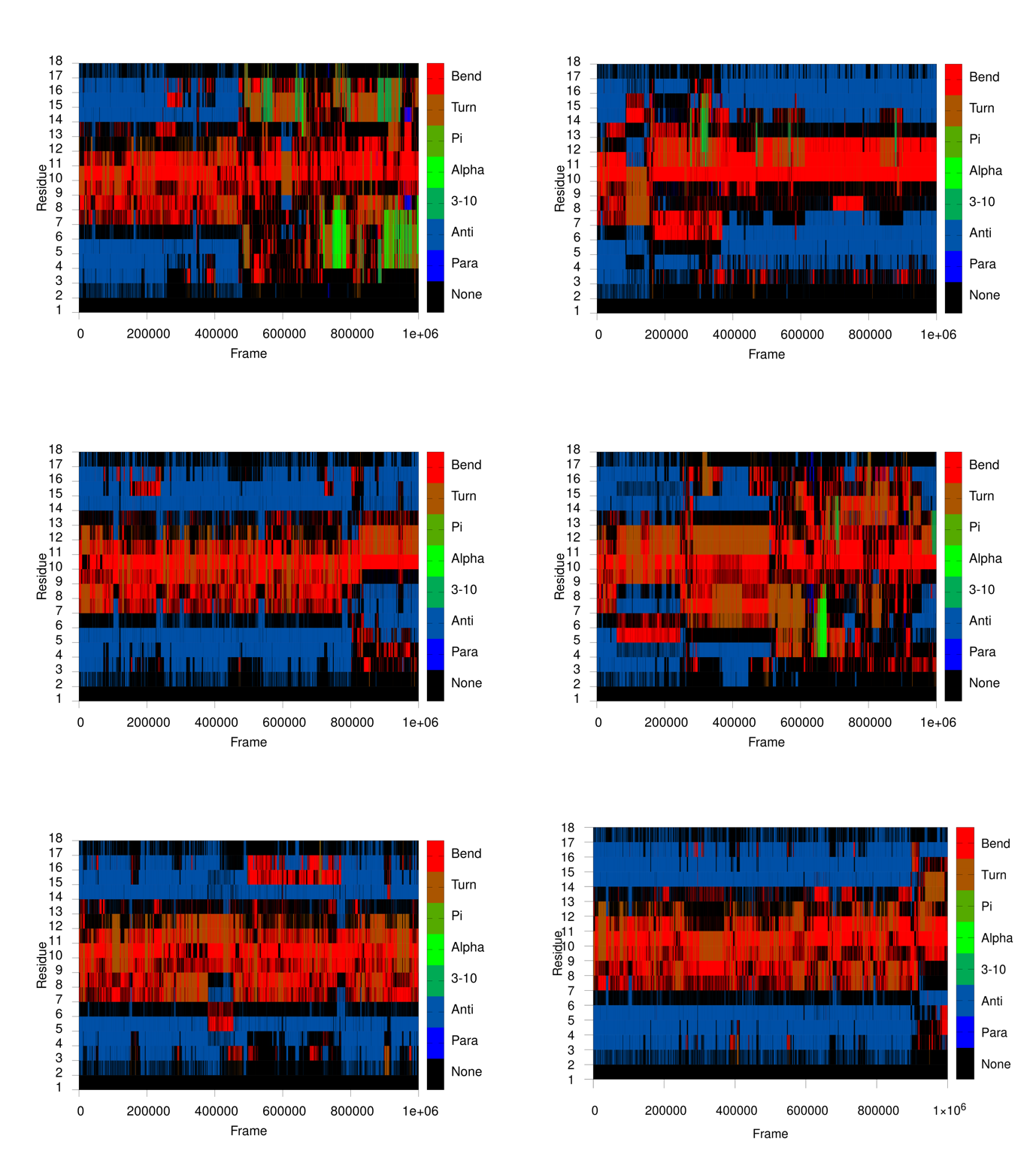

Figure S3. Evolution of secondary structure elements of HIV-1 peptide IIIB V3, wild-type β-hairpin at 77 °C during 1 μs MD simulation (six replicas). Colors denote certain element of secondary structure. Para - parallel β-sheet, Anti - antiparallel β-sheet, 3-10 – 3_10_ helix, Alpha - α-helix, Pi - π-helix, Turn - β-turn and Bend.

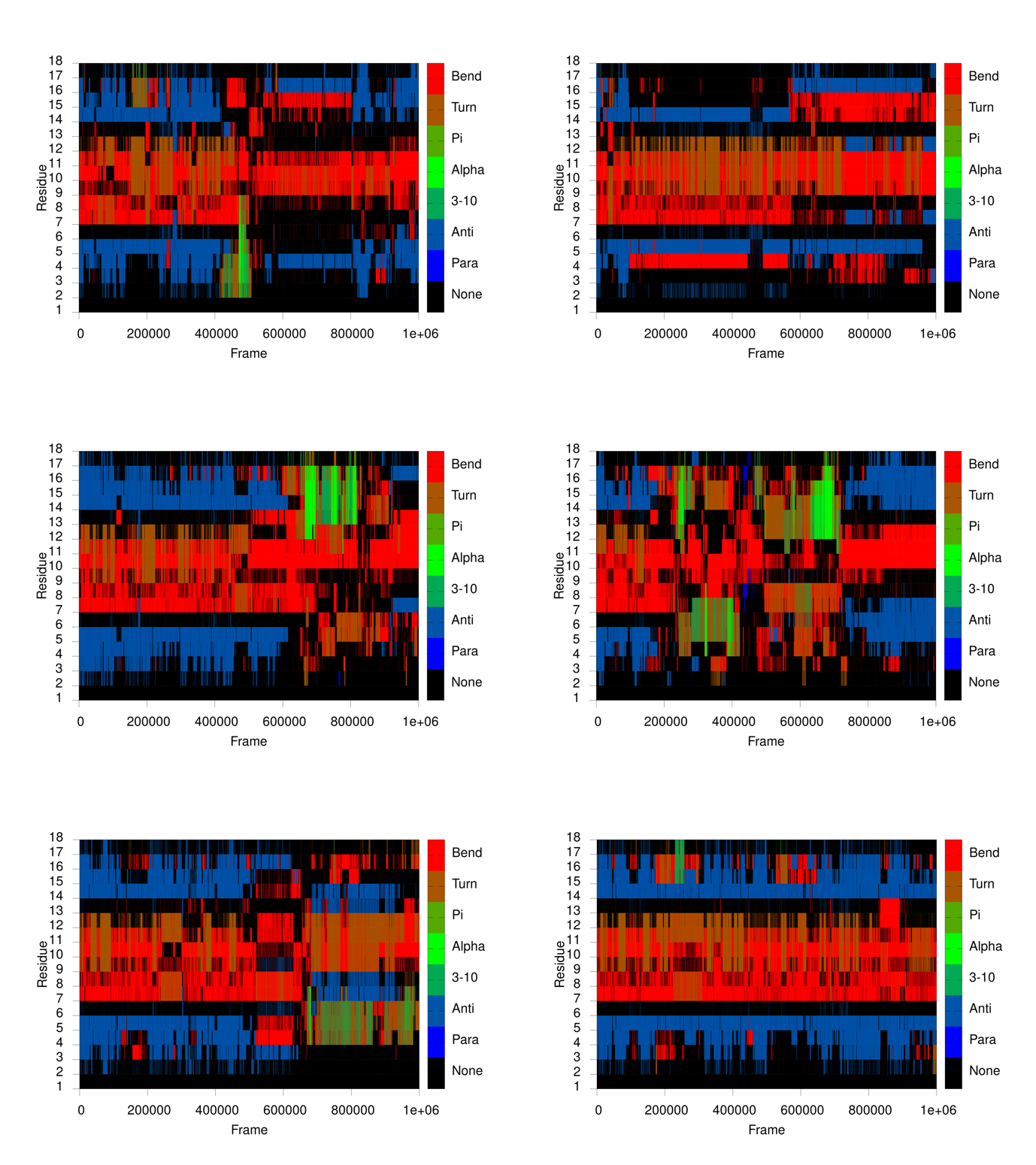

Figure S4. Evolution of secondary structure elements of HIV-1 peptide IIIB V3, β-hairpin with three alanines instead of isoleucines at 27 °C during 1 μs MD simulation (six replicas). Colors denote certain element of secondary structure. Para - parallel β-sheet, Anti - antiparallel β-sheet, 3-10 – 3_10_ helix, Alpha - α-helix, Pi - π-helix, Turn - β-turn and Bend.

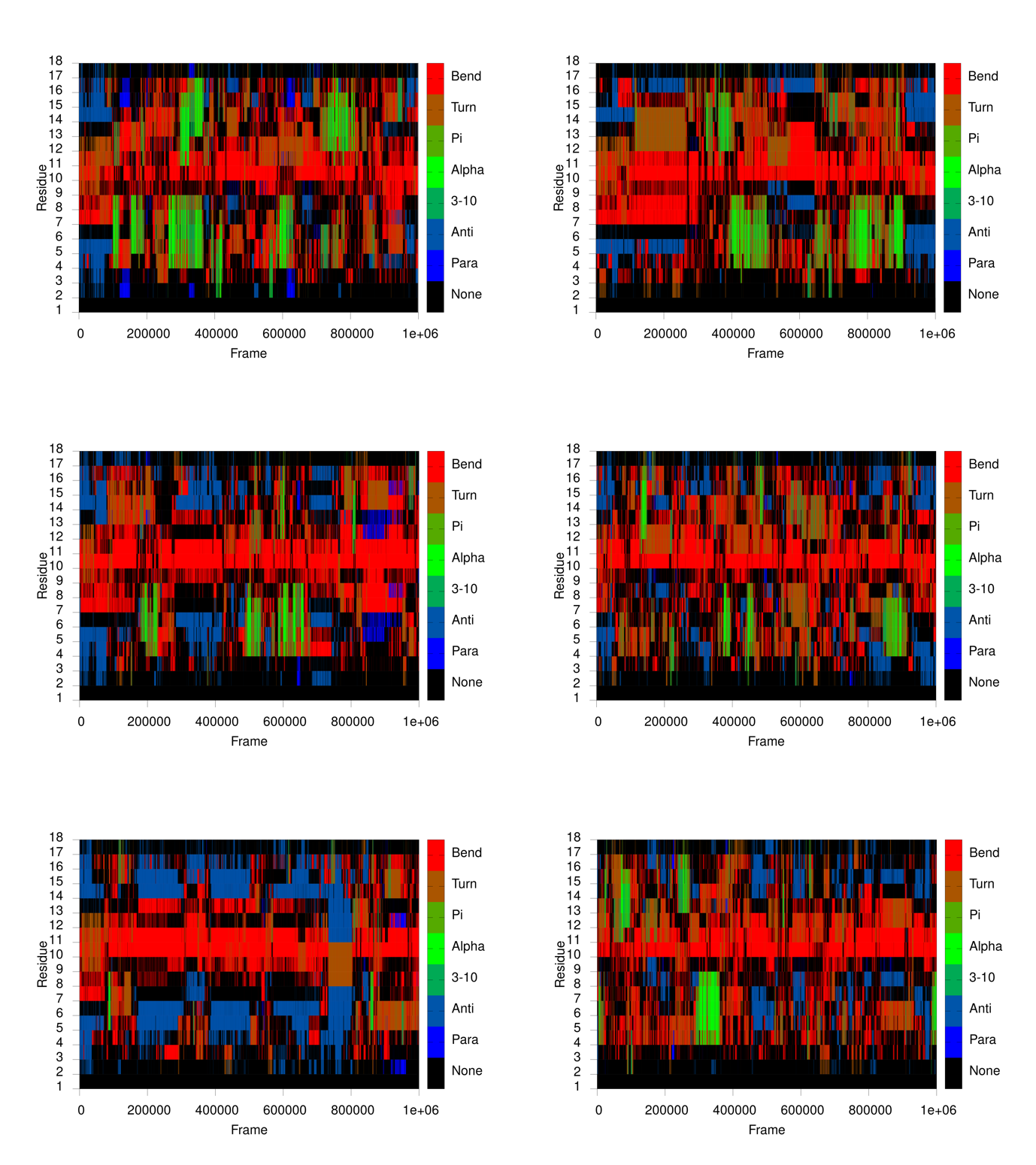

Figure S5. Evolution of secondary structure elements of HIV-1 peptide IIIB V3, β-hairpin with three alanines instead of isoleucines at 77 °C during 1 μs MD simulation (six replicas). Colors denote certain element of secondary structure. Para - parallel β-sheet, Anti - antiparallel β-sheet, 3-10 – 3_10_ helix, Alpha - α-helix, Pi - π-helix, Turn - β-turn and Bend.

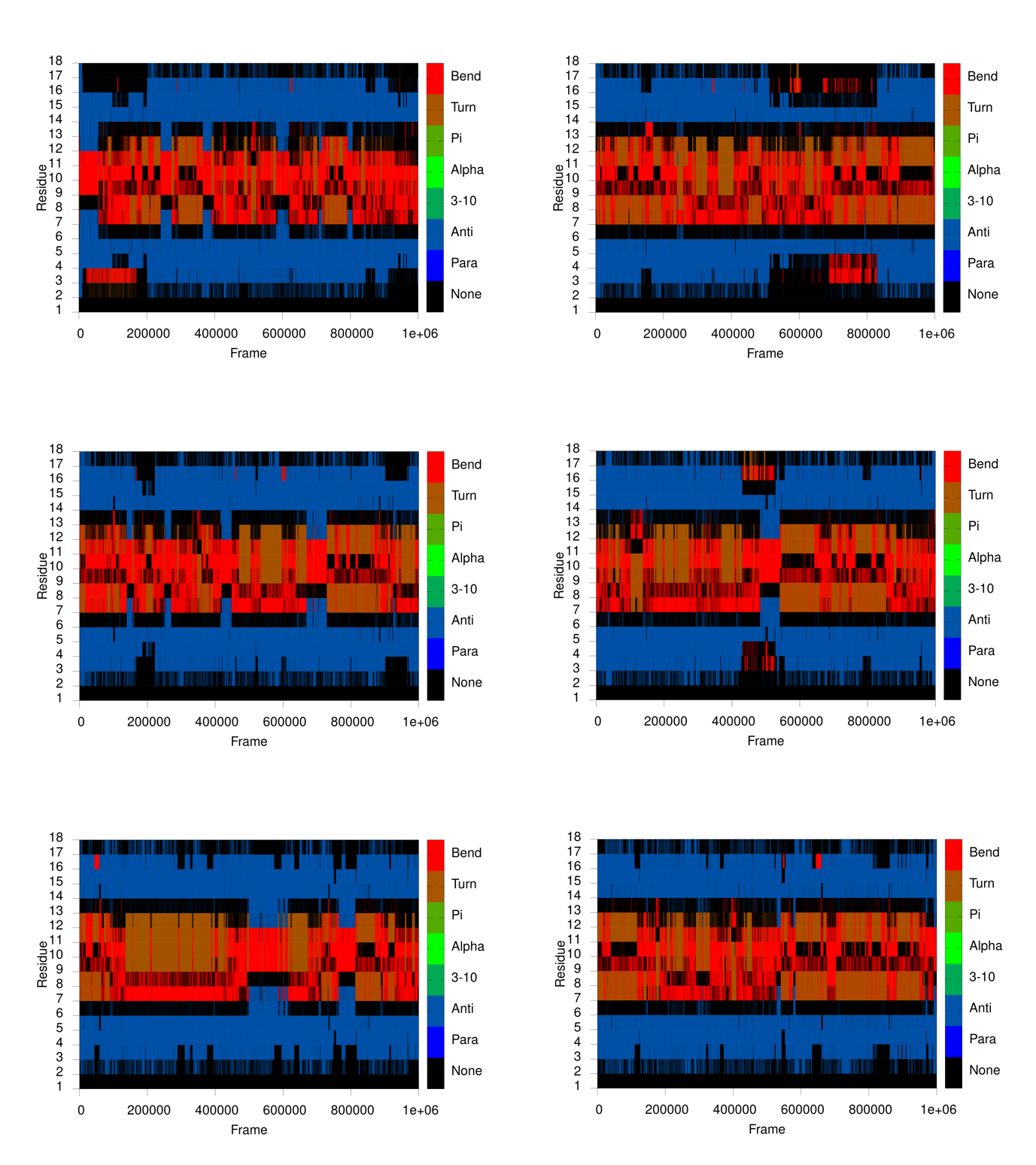

Figure S6. Evolution of secondary structure elements of HIV-1 peptide IIIB V3, β-hairpin with three valines instead of isoleucines at 27 °C during 1 μs MD simulation (six replicas). Colors denote certain element of secondary structure. Para - parallel β-sheet, Anti - antiparallel β-sheet, 3-10 – 3_10_ helix, Alpha - α-helix, Pi - π-helix, Turn - β-turn and Bend.

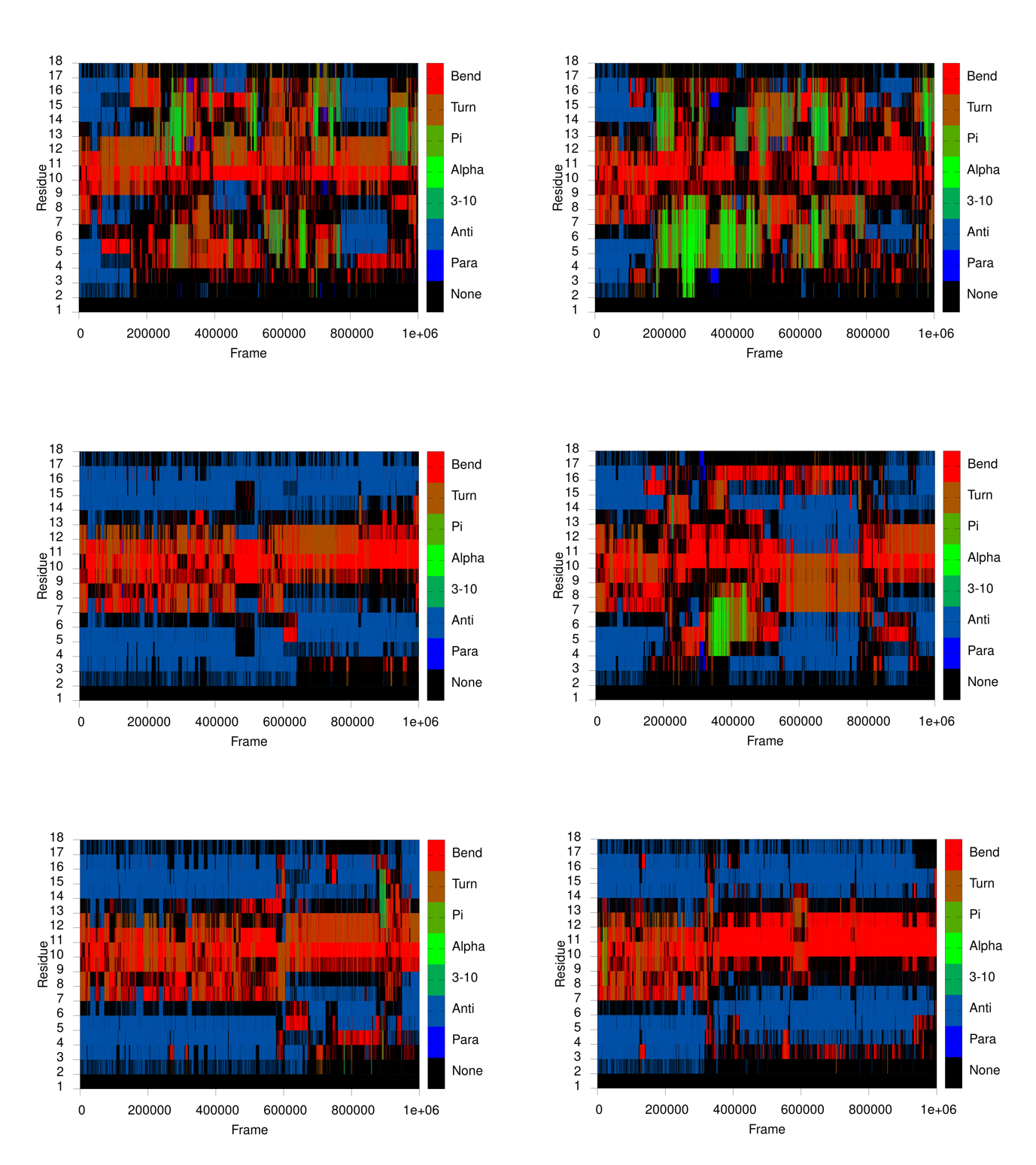

Figure S7. Evolution of secondary structure elements of HIV-1 peptide IIIB V3, β-hairpin with three valines instead of isoleucines at 77 °C during 1 μs MD simulation (six replicas). Colors denote certain element of secondary structure. Para - parallel β-sheet, Anti - antiparallel β-sheet, 3-10 – 3_10_ helix, Alpha - α-helix, Pi - π-helix, Turn - β-turn and Bend.

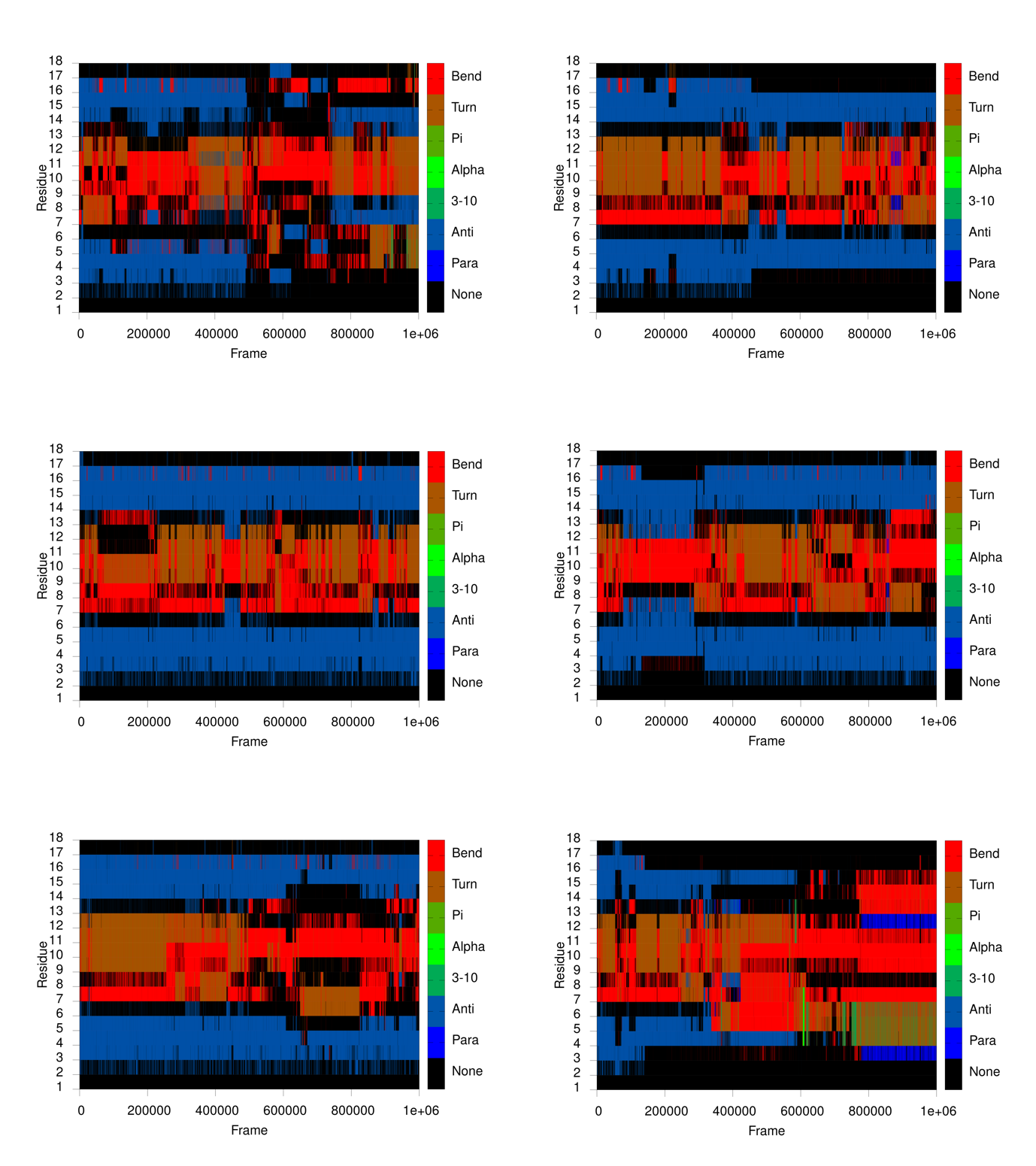

Figure S8. Evolution of secondary structure elements of HIV-1 peptide IIIB V3, β-hairpin with three norvalines instead of isoleucines at 27 °C during 1 μs MD simulation (six replicas). Colors denote certain element of secondary structure. Para - parallel β-sheet, Anti - antiparallel β-sheet, 3-10 – 3_10_ helix, Alpha - α-helix, Pi - π-helix, Turn - β-turn and Bend.

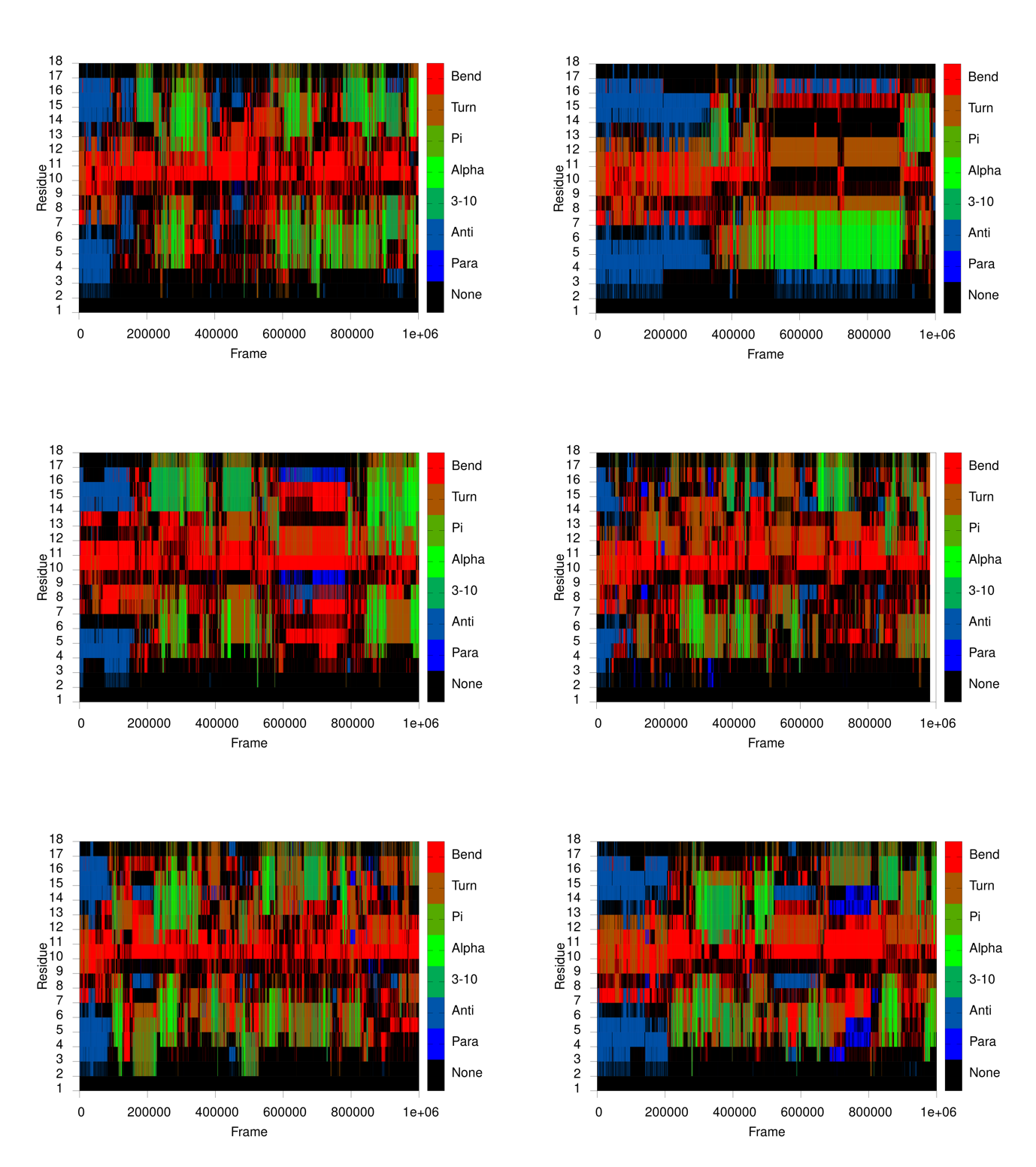

Figure S9. Evolution of secondary structure elements of HIV-1 peptide IIIB V3, β-hairpin with three norvalines instead of isoleucines at 77 °C during 1 μs MD simulation (six replicas). Colors denote certain element of secondary structure. Para - parallel β-sheet, Anti - antiparallel β-sheet, 3-10 – 3_10_ helix, Alpha - α-helix, Pi - π-helix, Turn - β-turn and Bend.

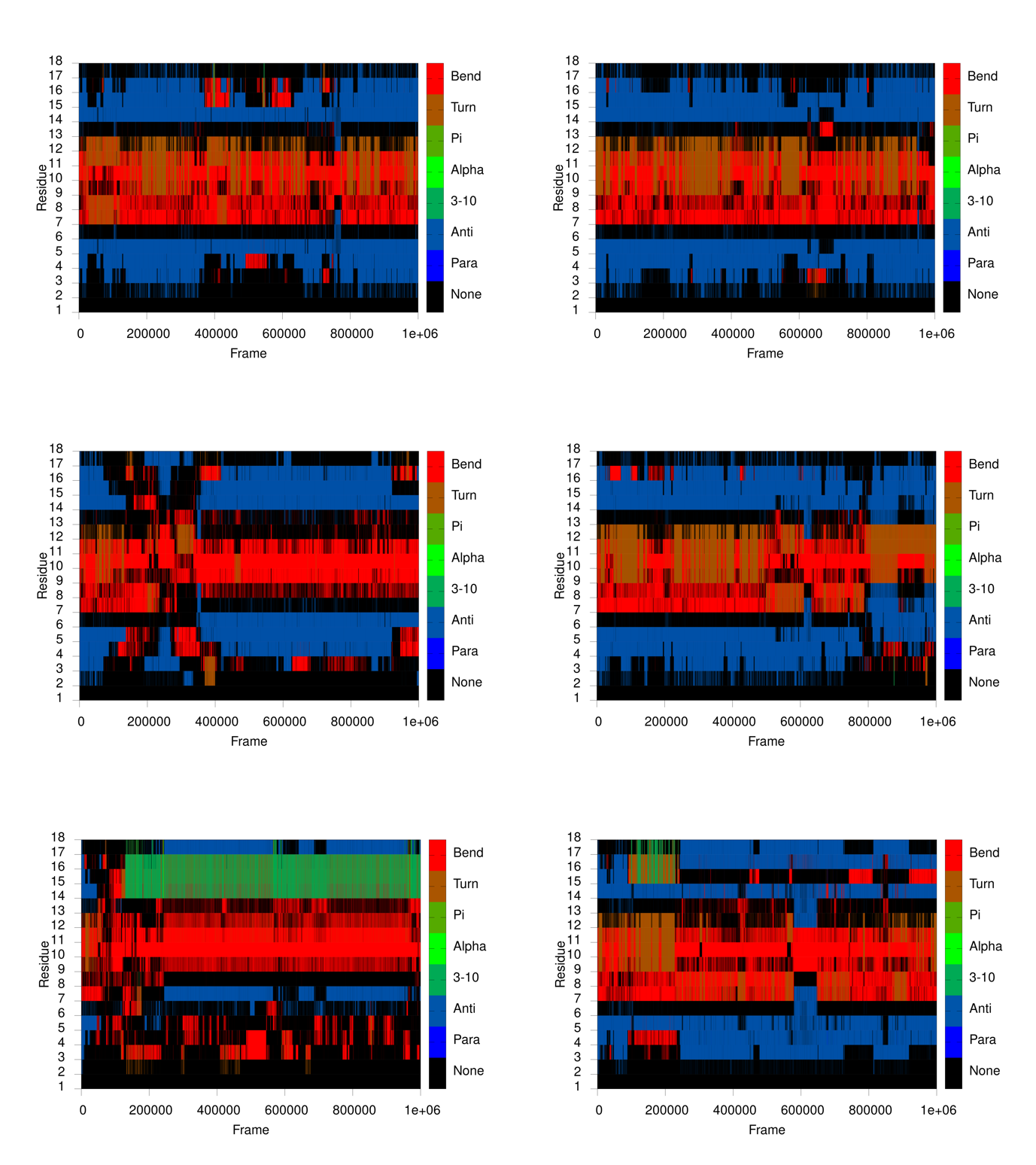

Figure S10. Evolution of secondary structure elements of HIV-1 peptide IIIB V3, β-hairpin with three leucines instead of isoleucines at 27 °C during 1 μs MD simulation (six replicas). Colors denote certain element of secondary structure. Para - parallel β-sheet, Anti - antiparallel β-sheet, 3-10 – 3_10_ helix, Alpha - α-helix, Pi - π-helix, Turn - β-turn and Bend.

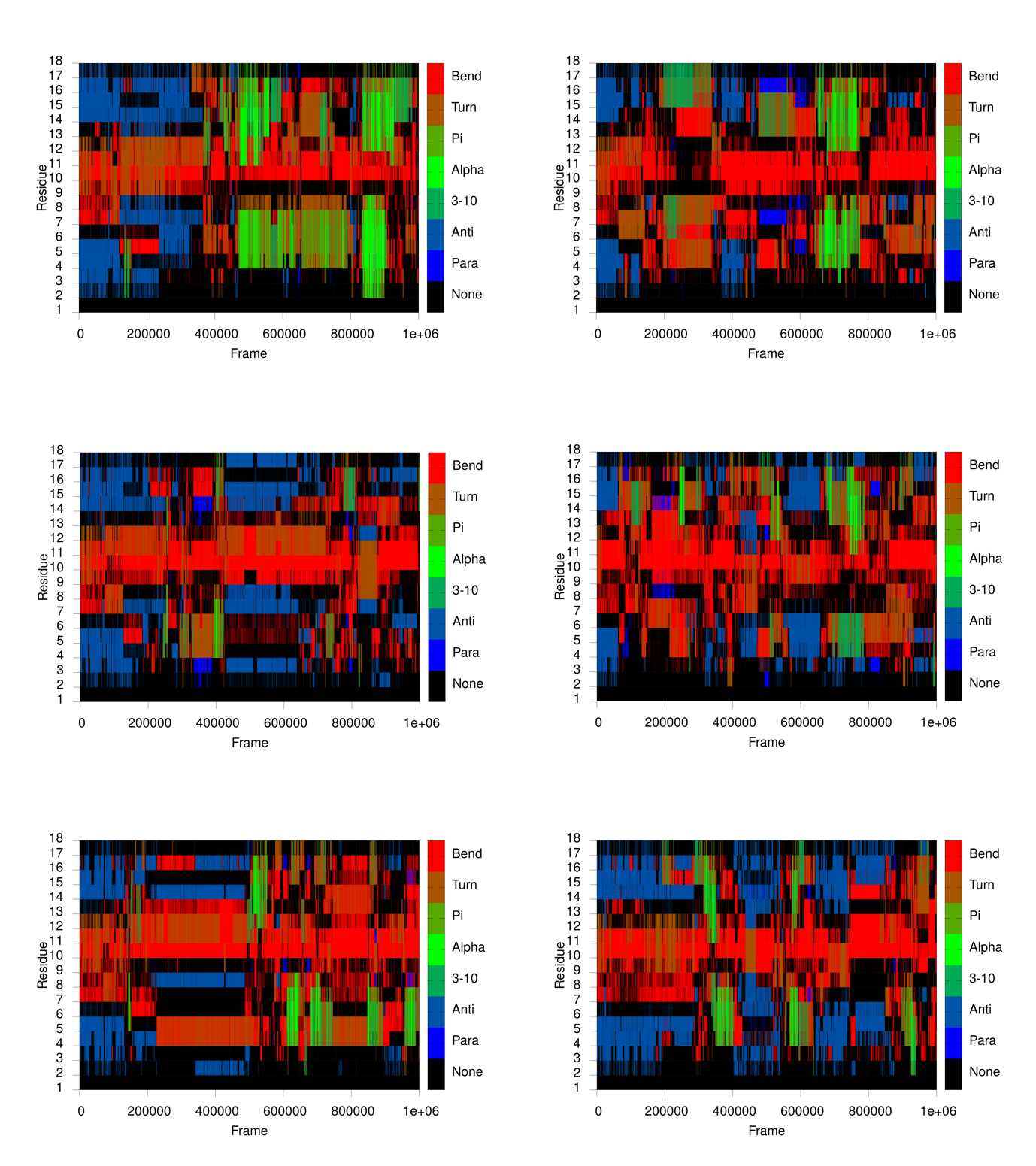

Figure S11. Evolution of secondary structure elements of HIV-1 peptide IIIB V3, β-hairpin with three leucines instead of isoleucines at 77 °C during 1 μs MD simulation (six replicas). Colors denote certain element of secondary structure. Para - parallel β-sheet, Anti - antiparallel β-sheet, 3-10 – 3_10_ helix, Alpha - α-helix, Pi - π-helix, Turn - β-turn and Bend.

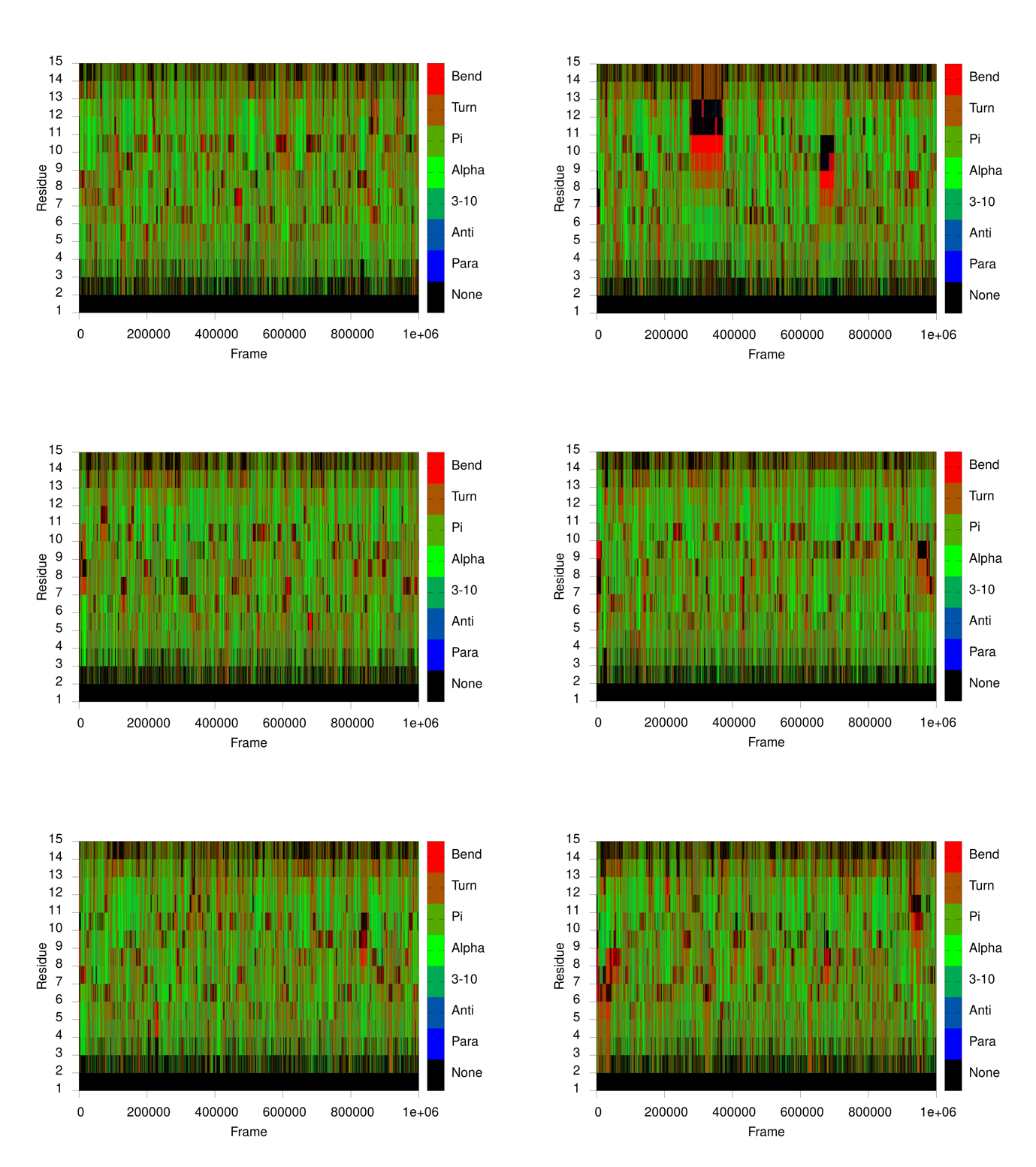

Figure S12. Evolution of secondary structure elements of polypeptide comprised of 13 norvalines with ACE and NME caps at 27 °C during 1 μs MD simulation (six replicas). Colors denote certain element of secondary structure. Para - parallel β-sheet, Anti - antiparallel β-sheet, 3-10 – 3_10_ helix, Alpha - α-helix, Pi - π-helix, Turn - β-turn and Bend.

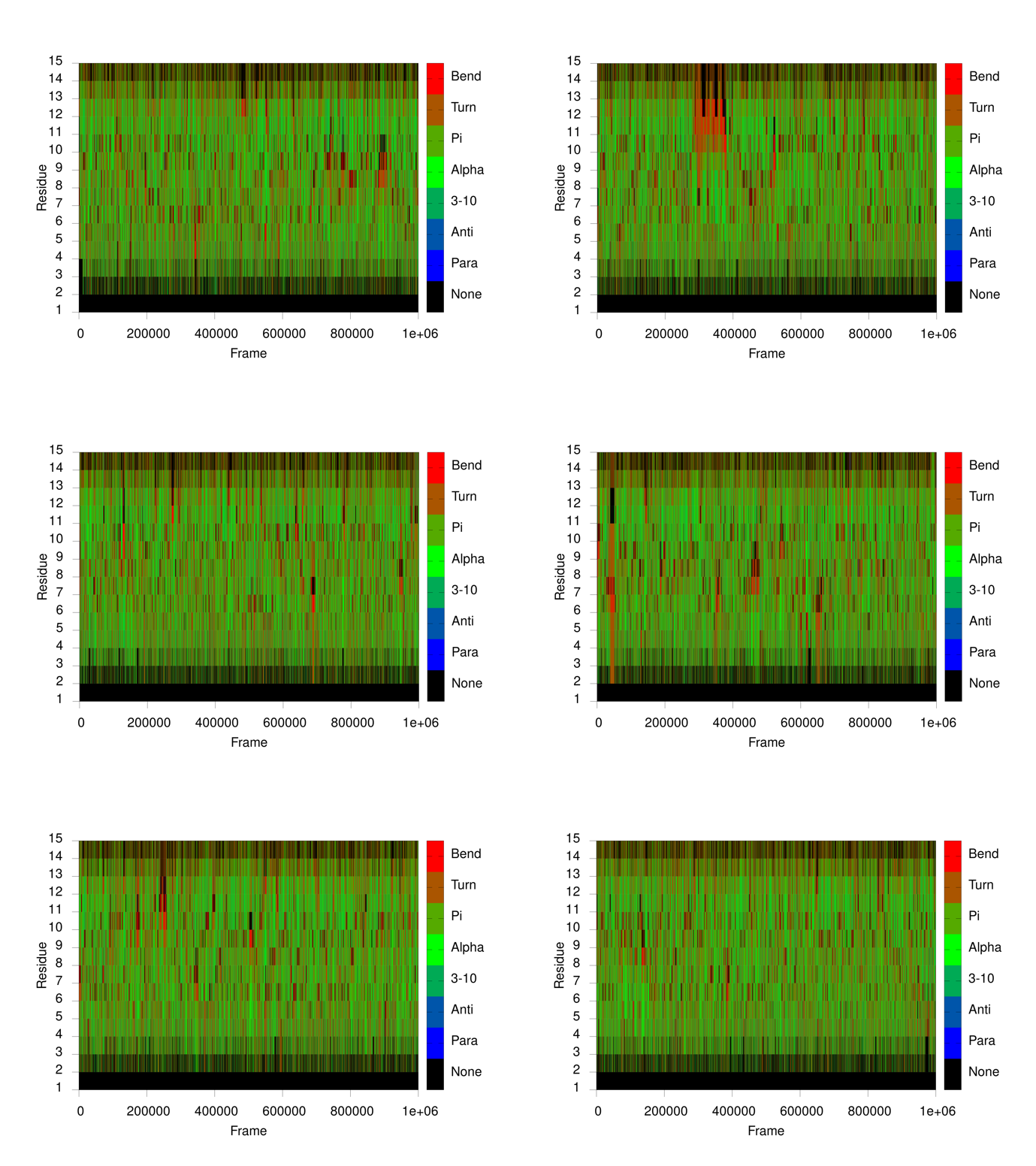

Figure S13. Evolution of secondary structure elements of polypeptide comprised of 13 norvalines with ACE and NME caps at 77 °C during 1 μs MD simulation (six replicas). Colors denote certain element of secondary structure. Para - parallel β-sheet, Anti - antiparallel β-sheet, 3-10 – 3_10_ helix, Alpha - α-helix, Pi - π-helix, Turn - β-turn and Bend.

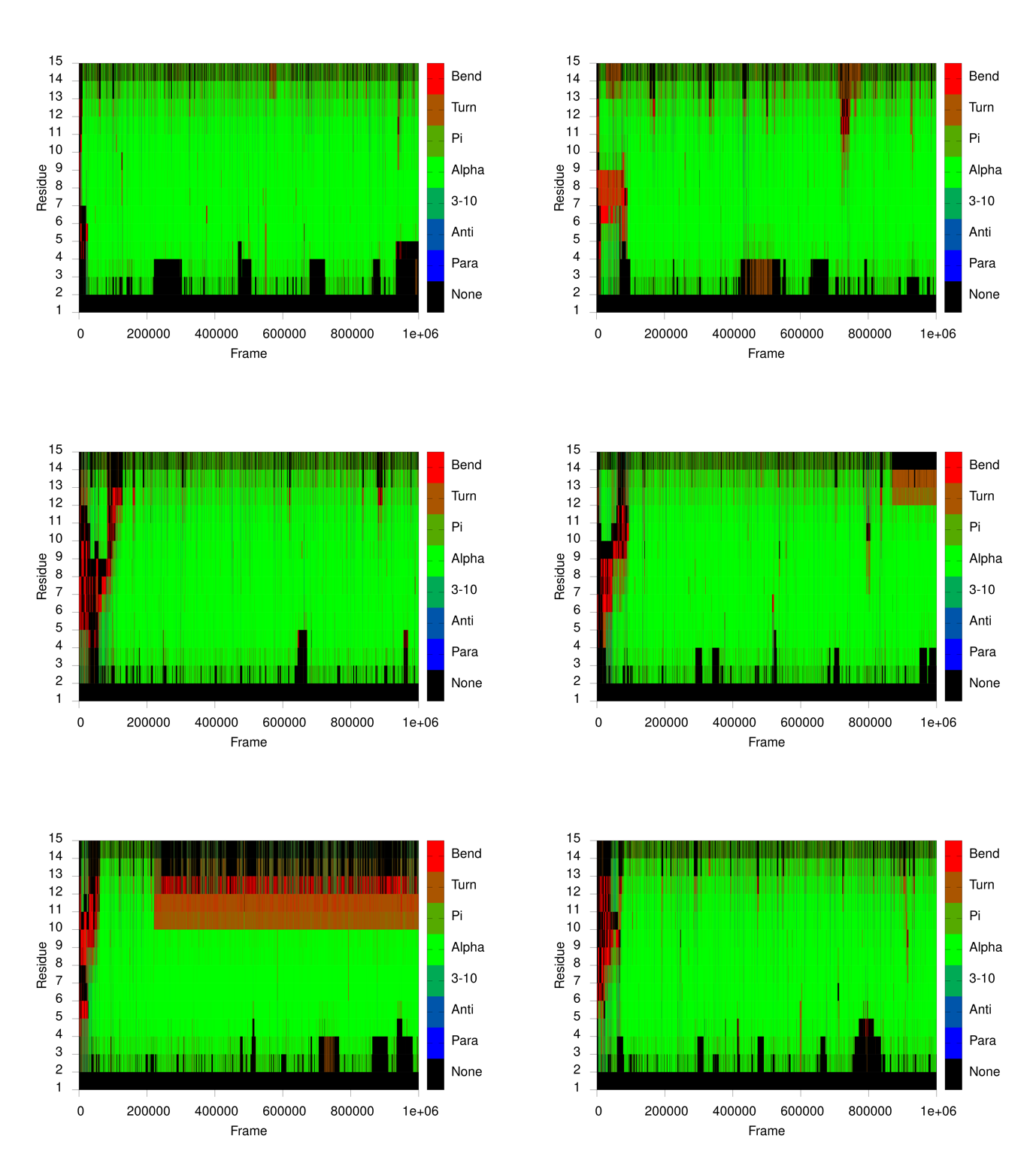

Figure S14. Evolution of secondary structure elements of polypeptide comprised of 13 leucines with ACE and NME caps at 27 °C during 1 μs MD simulation (six replicas). Colors denote certain element of secondary structure. Para - parallel β-sheet, Anti - antiparallel β-sheet, 3-10 – 3_10_ helix, Alpha - α-helix, Pi - π-helix, Turn - β-turn and Bend.

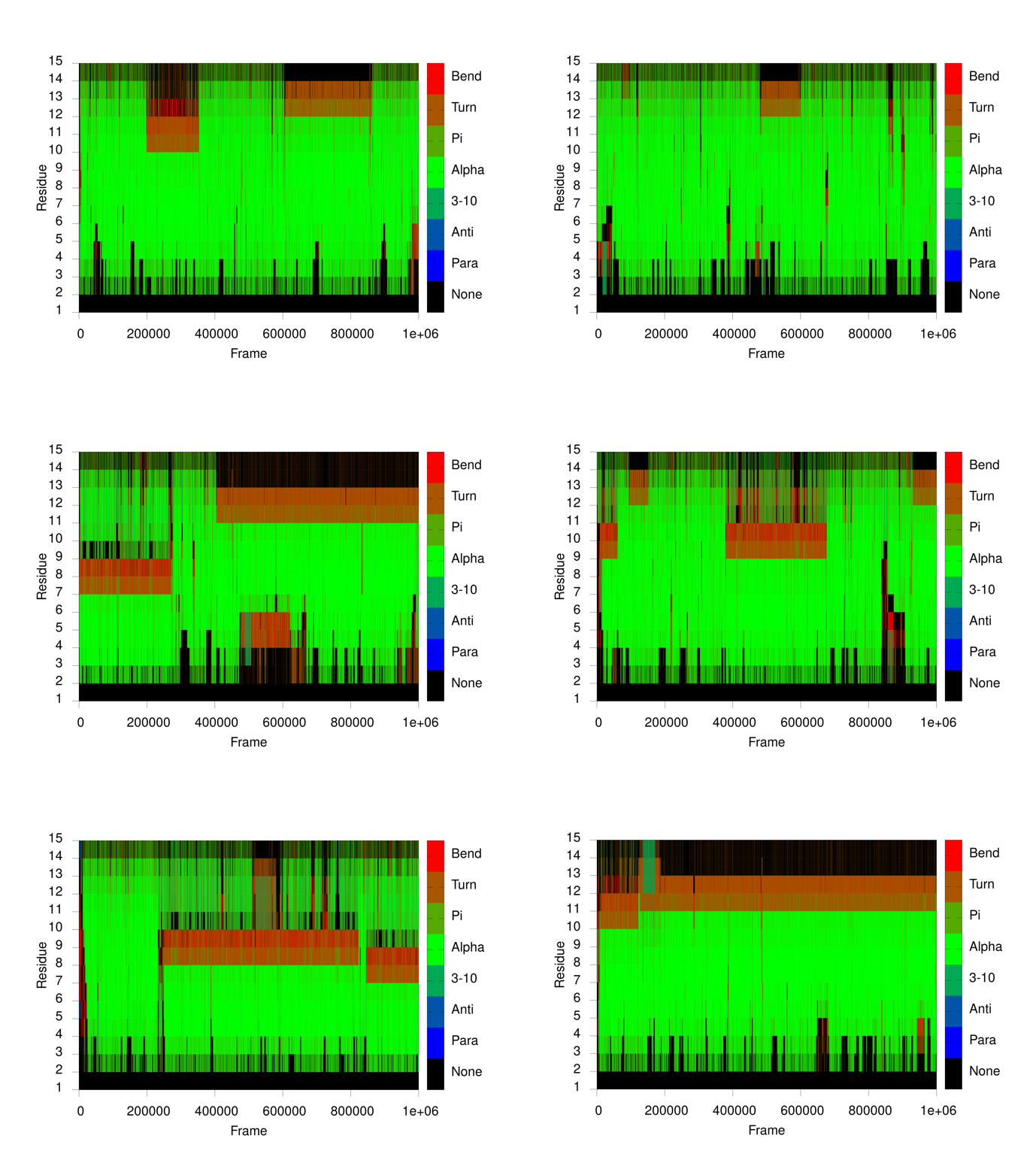

Figure S15. Evolution of secondary structure elements of polypeptide comprised of 13 leucines with ACE and NME caps at 77 °C during 1 μs MD simulation (six replicas). Colors denote certain element of secondary structure. Para - parallel β-sheet, Anti - antiparallel β-sheet, 3-10 – 3_10_ helix, Alpha - α-helix, Pi - π-helix, Turn - β-turn and Bend.
